# The Viability Gambit: An Optimized Sterilisation Protocol for Industrial Hemp (*Cannabis sativa* L.) Seeds

**DOI:** 10.64898/2026.06.25.732887

**Authors:** Rachna Gowlikar, Grace Pender, Joanna Kacprzyk, Alice Destailleur, Aditya Nayak, Rainer Melzer, Susanne Schilling

## Abstract

Hemp (*Cannabis sativa* L.) is an increasingly important crop with applications spanning fibre, seed oil and bioactive cannabinoid production, yet the development of reliable tissue culture systems for this species remains a significant challenge. The establishment of axenic seedling cultures is a prerequisite for hypocotyl-based regeneration and future genetic transformation pipelines, but hemp seeds harbour diverse endophytic microbial communities that frequently overwhelm standard surface sterilisation protocols.

Here, we present a systematic comparative evaluation of seed sterilisation strategies across seven industrial hemp accessions, examining the effects of sterilisation chemistry, seed provenance and accession identity on both contamination outcome and the subsequent morphogenic competence of hypocotyl explants. Across all treatments and accessions, in-house glasshouse-harvested seeds achieved higher sterility rates than commercially sourced material regardless of protocol applied. This provenance effect, combined with considerable batch-to-batch variation within seed sources, indicates that contamination load is a primary determinant of successful hemp seed sterilisation. Among the sterilisation treatments evaluated, a baseline of 75% ethanol combined with sequential 1% hydrogen peroxide incubation performed consistently well for low-load seed batches, while supplementation with Plant Preservative Mixture (PPM^TM^) might be necessary to achieve acceptable rates of non-contaminated seedlings from high contamination load batches. Beyond their effect on contamination, sterilisation treatments influenced the morphogenic fate of hypocotyl explants independently of sterility outcomes. Notably, seedling treatment with the Prochloraz-based fungicide Octave promoted shoot and root co-regeneration in the absence of exogenous plant growth regulators in some cases. Hormone-free organogenesis from hypocotyl explants was achievable across multiple hemp accessions, demonstrating that this developmental capacity is broadly distributed within hemp, though its frequency and consistency varied with accession identity and protocol conditions.

Together, these findings provide a practical framework for axenic hemp seed culture that can be used as starting point requiring local adaptation based on seed source, batch history and the intended downstream application.

## Introduction

Hemp (*Cannabis sativa* L.) is a highly versatile crop with a broad range of industrial, agricultural and pharmacological applications, including the production of fibre, seed oil and bioactive cannabinoids such as cannabidiol (CBD) (Andre et al., 2016; Schilling et al., 2023; Small, 2015). Following decades of regulatory restriction, hemp cultivation has undergone a global resurgence and is now practiced across more than 30 countries. This expansion has created a growing demand for scalable, genetically consistent planting material and has renewed interest in developing robust tissue culture systems for this species (Chandra et al., 2017).

Tissue culture and plant regeneration are foundational tools in modern crop research and breeding. The ability to regenerate whole plants from isolated explants is a prerequisite for genetic transformation, enabling the stable introduction of defined traits and the functional characterisation of genes of interest. In many crop species, efficient transformation and regeneration pipelines have accelerated both basic research and the development of improved varieties (Hesami et al., 2021). In hemp, however, these tools remain poorly developed. Efficient and reproducible regeneration protocols are lacking, and genetic transformation of *C. sativa* has not yet been reliably achieved, representing a significant bottleneck for both fundamental research and applied breeding programmes.

Hypocotyl explants are among the most promising target tissues for hemp tissue culture and potential transformation, as they consist of rapidly dividing cells with broad developmental plasticity and have been shown to undergo organogenesis under hormone-free conditions in some accessions (Galán-Ávila et al., 2020; Hesami et al., 2021). However, the utility of hypocotyl-based regeneration systems depends entirely on the successful axenic establishment of seedlings from surface-sterilized seeds. This upstream step is non-trivial in hemp. Seeds harbour a taxonomically diverse endophytic microbiome, including bacterial genera such as *Pseudomonas*, *Pantoea* and *Bacillus*, and fungal endophytes from genera including *Alternaria*, *Cladosporium* and *Aureobasidium*, which are deeply embedded within seed tissues and resistant to standard surface sterilisation (Punja et al., 2018). *In vitro* cultures are additionally susceptible to fast-growing opportunistic pathogens common in hemp production environments, including *Botrytis cinerea* and *Penicillium* spp. (Punja et al., 2019). The combination of resident endophytes and surface contaminants makes sterilisation protocols developed for model species such as *Arabidopsis thaliana* or *Nicotiana tabacum* frequently inadequate when applied to hemp (Srivastava, 2024).

A further complication is the substantial genetic and phenotypic diversity among industrial hemp cultivars (Kavanagh et al., 2025; Trubanová et al., 2026; Vergara et al., 2017). Cultivars are highly heterozygous and vary considerably in seed morphology and coat properties, which likely influences both the depth of microbial colonisation and the response to chemical sterilants (Hesami et al., 2021). Seed provenance adds another layer of variability: field-harvested commercial seeds are exposed to diverse soil and airborne microbial communities during development, harvest and storage, whereas seeds produced under controlled glasshouse conditions carry a lower and more predictable contamination burden. These sources of variation are rarely accounted for systematically in published sterilisation protocols, which are typically reported for a single cultivar and seed source without broader validation.

Here, we present a comparative analysis of seed surface sterilisation across seven industrial hemp accessions, evaluating standalone chemical treatments including H₂O₂ and NaOCl at varying concentrations alongside combinatorial protocols using a standard ethanol and sodium hypochlorite baseline supplemented with the Prochloraz-based fungicide Octave and Plant Preservative Mixture^TM^ (PPM^TM^). Critically, this study extends beyond contamination metrics alone to examine how sterilisation treatment, seed provenance and accession identity jointly influence the morphogenic competence of hypocotyl explants in subsequent hormone-free culture. The aim is to provide a practical and honestly characterised framework for axenic hemp seed culture that acknowledges the batch- and context-dependent nature of sterilisation outcomes and supports the development of more robust regeneration systems for this increasingly important crop.

## Materials and Methods

### Plant material and cultivation

Hemp seeds from seven cultivars (’Felina 32’, ’Ferimon’, ’Santhica 27’, ’Kompolti’, ’USO31’, ’Futura 75’ and ’Fedora 17’) were obtained from two sources: commercially certified field-harvested seed lots (HEMPOINT, Czech Republic) and in-house produced seed stocks. In-house seeds of were generated from plants cultivated under controlled glasshouse conditions at University College Dublin, maintained under glasshouse conditions following (Schilling et al., 2023). All seeds were stored at 4°C in darkness until use. Hemp explants on were grown on ½ MS medium in a plant cultivation room under red/blue LED lights at a constant temperature of 25°C under a 16-hour photoperiod.

### Surface sterilisation

All sterilisation protocols began with a common ethanol baseline. Seeds were agitated in 75% ethanol for three minutes and rinsed three times with sterile distilled water. For water, vinegar (1%) and salt (1%) treatment, seeds were agitated in the respective solution for 15 minutes. For sodium hypochlorite treatments seeds were agitated in 20% or 50% commercial bleach (approximately 1% and 3% NaOCl) at 72 rpm for 20 minutes, followed by four to five rinses with sterile distilled water. In the hydrogen peroxide treatment, seeds were soaked in 1% or 3% H₂O₂ in darkness at 25°C in sequential incubations of 24, 48, 24 hours with sterile water washes and fresh H₂O₂ solution between incubations. For the Octave treatment, seeds were agitated in a Prochloraz-based fungicide solution (Octave, Bayer CropScience) at 180 rpm for two hours following the baseline, rinsed four to five times with sterile distilled water, then incubated in sterile water in darkness for 48 hours before transfer to growth medium.

Plant Preservative Mixture^TM^ (PPM™, Plant Cell Technology), a broad-spectrum biocide for plant tissue culture, was not applied as a seed treatment but incorporated directly into ½ Murashige and Skoog (MS) basal salt growth medium at the manufacturer’s recommended concentration (0.1%), providing continuous biocidal activity throughout the culture period. These treatments were applied individually and in combination.

### *In vitro* regeneration from hemp hypocotyl explants

Following sterilisation, hemp seeds were transferred to either agar-solidified ½ MS medium. For *in vitro* culture, seeds were plated on ½ MS medium. Sterile seedling yield was calculated as the proportion of plated seeds producing uncontaminated, viable seedlings. Hypocotyl explants were excised from sterile seedlings 7-14 days after germination and transferred to hormone-free ½ MS medium (Galán-Ávila et al., 2021). Explants were scored for regeneration outcome after 14 days, with responses categorized as shoot-only regeneration, root-only regeneration, combined shoot and root regeneration, or no regeneration.

### Statistical analysis and data visualisation

All statistical analyses and figures were generated in R (version 2026.04.0+526; R Core Team, 2026). Sterile seedling yield and explant regeneration outcome data were analysed using binomial generalised linear models (GLM), with pairwise comparisons performed using Tukey’s HSD post-hoc test implemented in the R emmeans package (cran.r-project.org/package=emmeans). Differences in morphogenic outcome distribution across accessions were assessed using a Chi-squared test of independence with simulated p-values. The overall effect of seed provenance on sterility rate was tested using a two-sample t-test. Figures were generated using the ggplot2 package (Wickham, 2016).

## Results

### Ethanol combined with hydrogen peroxide or sodium hypochlorite are effective for sterilising hemp seeds

To establish protocols suitable for routine axenic tissue culture for hemp, seeds of different hemp cultivars underwent a variety of treatments with different degrees of stringency and were subsequently germinated on agar plates and assessed (**Figure 1a-c**). Water-only controls and salt- or vinegar-supplemented seed treatments produced near-universal contamination or germination failure across all accessions, confirming that more stringent seed treatment is a prerequisite for successful establishment tissue cultures (**Figure S1**). Further, treatment using 75% ethanol combined with 1% H₂O₂ produced contamination-free seeds with an acceptable seedling viability of around 25 – 75% (**Figure 1d, Figure S1**). This treatment was therefore carried forward as a baseline for subsequent experiments.

**Figure 1.**
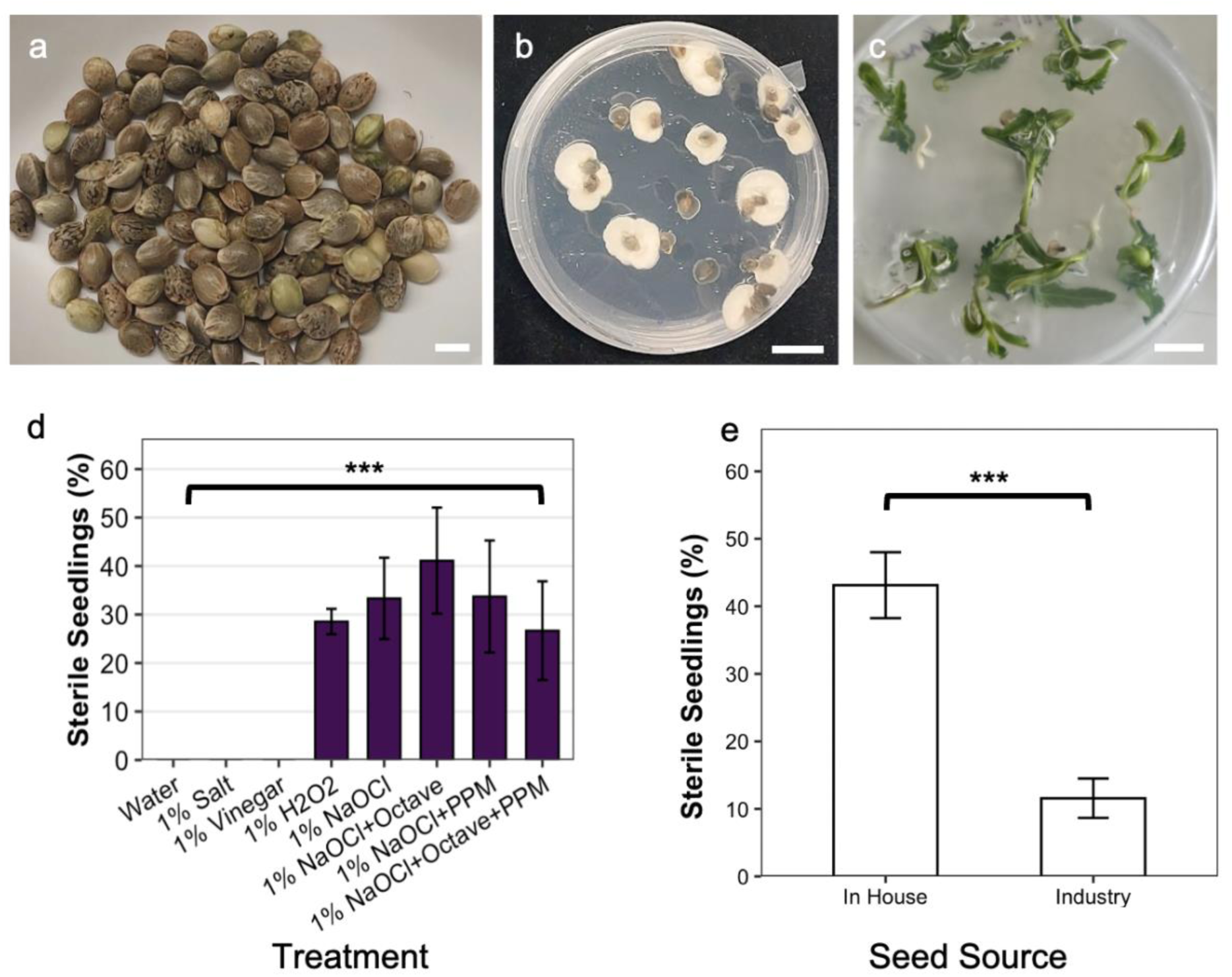
Effect of varying seed disinfection protocols on hemp seedling sterility. Hemp seeds were treated with different sterilisation protocols and were scored if they were contaminated (b) or sterile (c). The mean percentage of sterile seedlings varied depending on the sterilisation protocol used (d), with significant differences between water (n=135), salt (n=20) or vinegar (n=20) treated plants and all other sterilisation methods, including 1%H2O2 (n=289), 1% NaOCl (n=90), 1% NaOCl combined with Octave (n=90), 1% NaOCl combined with PPM^TM^ (n=90) and 1% NaOCl combined with Octave and PPM^TM^ (n=90) (Tukey’s HSD, ***, p<0.001). Overall, seedlings derived from seeds that were grown in-house (n=3441) showed significantly higher sterility rates as compared to commercially sourced seeds (’industry’, n=1886) over a variety of different sterilisation treatments (e, ***, p<0.001) (Table S1). Scale bar 5 mm (a) and 10 mm (b,c). Error bars indicate standard error (d, e).

Next, four NaOCl-based protocols were compared to assess whether supplementary biocidal agents improve sterile seedling yield (**Figure 1d, Figure S2**). In addition to 1 % NaOCl alone we supplemented NaOCl with the fungicide Octave and Plant Preservative Mixture (PPM™). Here, the yield of non-contaminated seedlings was significantly higher than in water, salt and vinegar treatments (Tukey’s HSD, p<0.001), but broadly similar across 1% NaOCl alone, NaOCl combined with Octave, NaOCl combined with PPM™, and NaOCl combined with both Octave and PPM™, with no statistically significant differences detected among the latter treatments (**Figure 1d**). In commercially sourced seeds, NaOCl combined with PPM™ performed significantly better than all other NaOCl based treatments (**Figure S2**).

While all experiments outlined so far used seeds produced locally in our greenhouse or growth rooms (termed in-house seeds henceforth), we also tested H₂O₂ and NaOCl based protocols on commercially sourced seeds. Commercially sourced seeds consistently showed a low percentage of sterile seedling across all four treatments, with means remaining below 10% regardless of the protocol applied (**Figure 1e, Table S1**). Neither Octave nor PPM meaningfully improved outcomes for commercially acquired seeds. The observation that in-house seeds produced a higher percentage of sterile seedlings when compared to industry sourced seeds persisted across all accessions and treatments in our screens. In-house seeds achieved roughly four times the sterile seedling rate of commercially sourced material across a variety of different hemp cultivars and treatments in our hands (Student’s t-test, p<0.001; **Figure 1e, Table S1**).

### Sterilisation treatments influence hemp hypocotyl regeneration outcomes

One important aspect of tissue culture is plant regeneration. For hemp, regeneration of plants from hypocotyls is one prominent approach. To test hypocotyl regeneration for hemp, hypocotyl explants recovered from sterilized seeds were transferred to hormone-free ½ MS medium and scored for regeneration outcome after 14 days. Notably, we observed bipolar regeneration from hypocotyl explants, with both shoot and roots developing from opposite sides of the same explant, referred further as root and shoot regeneration. Regeneration of only either shoot or root from hypocotyl explants was also frequently observed, termed either shoot only or root only, respectively.

Different seed sterilisation treatments were analysed for their effect on hypocotyl regeneration. Most hypocotyl explants generated from seeds of the cultivar ‘Felina 32’ (in-house) sterilized with 1% H_2_O_2_ or NaOCl died or regenerated only roots, with only a minor proportion producing shoots or shoots and roots (**Figure 2**). In contrast, explants from seeds sterilized with NaOCl combined with Octave as well as NaOCl combined with Octave and PPM showed a significantly higher proportion of shoot and root regeneration compared to sole NaOCl or H_2_O_2_ treatments (Fisher’s Exact Test, p<0.01). Additionally, explant mortality was significantly lower in explants originally sterilized with 1% NaOCl combined with PPM™ as compared to those treated with 1% NaOCl alone (Fisher’s Exact Test, p<0.001, **Figure 2**). In contrast, explants derived from seeds of the cultivar ‘Ferimon’ (commercial), regenerated shoots as well as roots best when initially sterilized with H_2_O_2_ without the addition of more stringent sterilisation agents (**Figure S3**).

**Figure 2.**
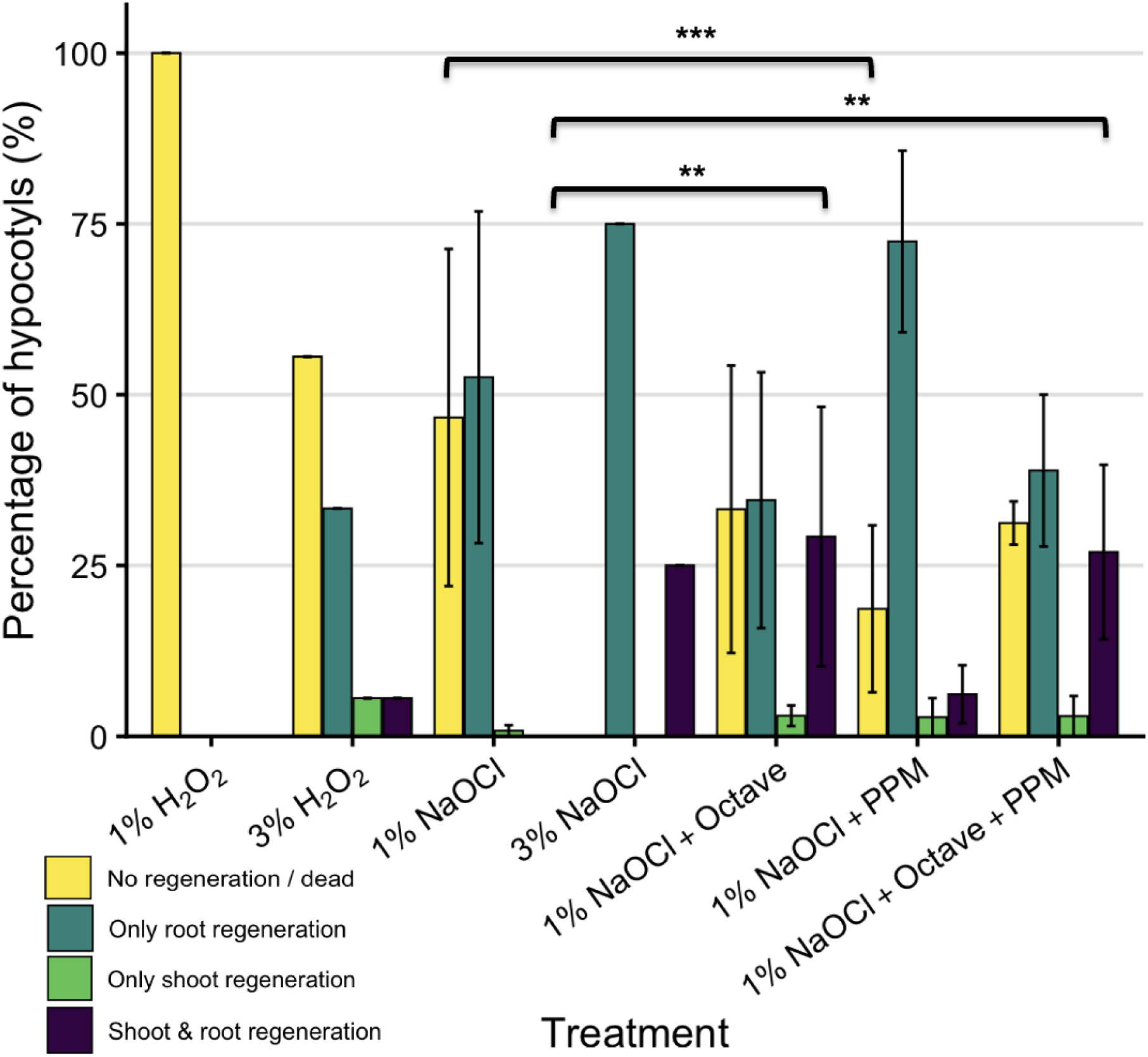
Effect of different sterilisation additives on the regeneration outcomes and viability of hemp hypocotyl explants. Explants derived from seedlings of the hemp cultivar ’Felina 32’ which underwent different sterilisation protocols (x-axis) were scored for their regeneration and categorized as no regeneration (yellow), only root regeneration (teal), only shoot regeneration (green) and shoot and root regeneration (dark blue). Selected significant differences are indicated with backets (Fishers exact test, **, p<0.01; ***, p<0.001).

### Regeneration capacity varies across hemp accessions and is influenced by seed provenance

Given the different responses of the two cultivars tested, we assessed the regeneration of hypocotyl explants across multiple accessions under a standardised sterilisation protocol to establish whether regeneration competence is broadly conserved or variable within hemp (**Figure 3**). Explants from four hemp cultivars were evaluated following sterilisation with 1% H₂O₂ (’Fedora 17’, ’Felina 32’, ’Kompolti’ and ’USO31’). Significant variation in morphogenic outcome distribution was detected among the accessions (Chi-squared test, χ^2^=27.71, df =9, p<0.01). Shoot and root co-regeneration was most frequent in ‘Fedora17’, with around a third of explants, while co-regeneration rate was significantly lower in ‘Kompolti’ and ‘Felina 32’ (Fishers exact test, p<0.05). However, in all cultivars around half of the total explants did not regenerate (**Figure 3**).

**Figure 3.**
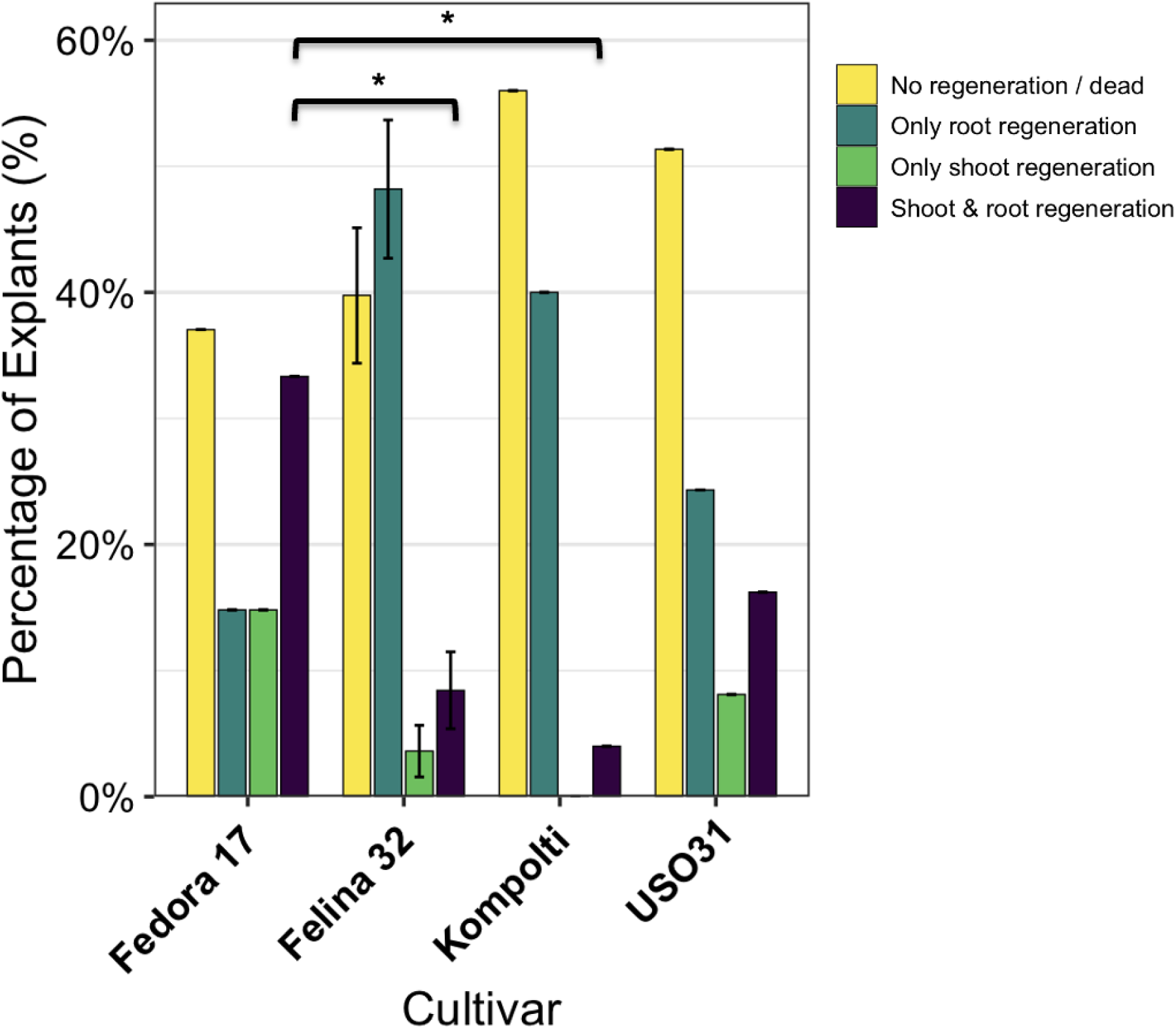
Regeneration outcomes across various hemp cultivars following 1% H₂O₂ surface sterilisation. Explants of seeds sterilized with 1% H_2_O_2_ of different hemp cultivars (’Fedora 17’ (n=27), ’Felina 32’ (n=83), ’Kompolti’ (n=25), ’USO 31’ (n=37)) were scored for their regeneration and categorized as no regeneration (yellow), only root regeneration (teal), only shoot regeneration (green) and shoot and root regeneration (dark blue). Selected significant differences are indicated with backets (Fishers exact test, *, p<0.05).

### Seed-to-seed regeneration via hypocotyl explants in hemp

To demonstrate that sterilized explants can complete the full regeneration cycle, the developmental progression from hypocotyl explant to fertile adult plant was documented (**Figure 4**). Sterile hypocotyl explants placed on hormone-free ½ MS medium developed shoots within two weeks (**Figure 4a-d**). Explants regenerated shoots and roots from the same hypocotyl in a polar way, with one end developing shoots and the other developing roots (**Figure 4e**) after approximately 3 weeks. Plantlets that established both roots and shoots were subsequently transferred to soil and grown to maturity in a growth room (**Figure 4f**). Plants derived from this protocol reached reproductive maturity and set seed (**Figure 4g, h**) which were successfully germinated and grown into a mature plant (not shown).

**Figure 4.**
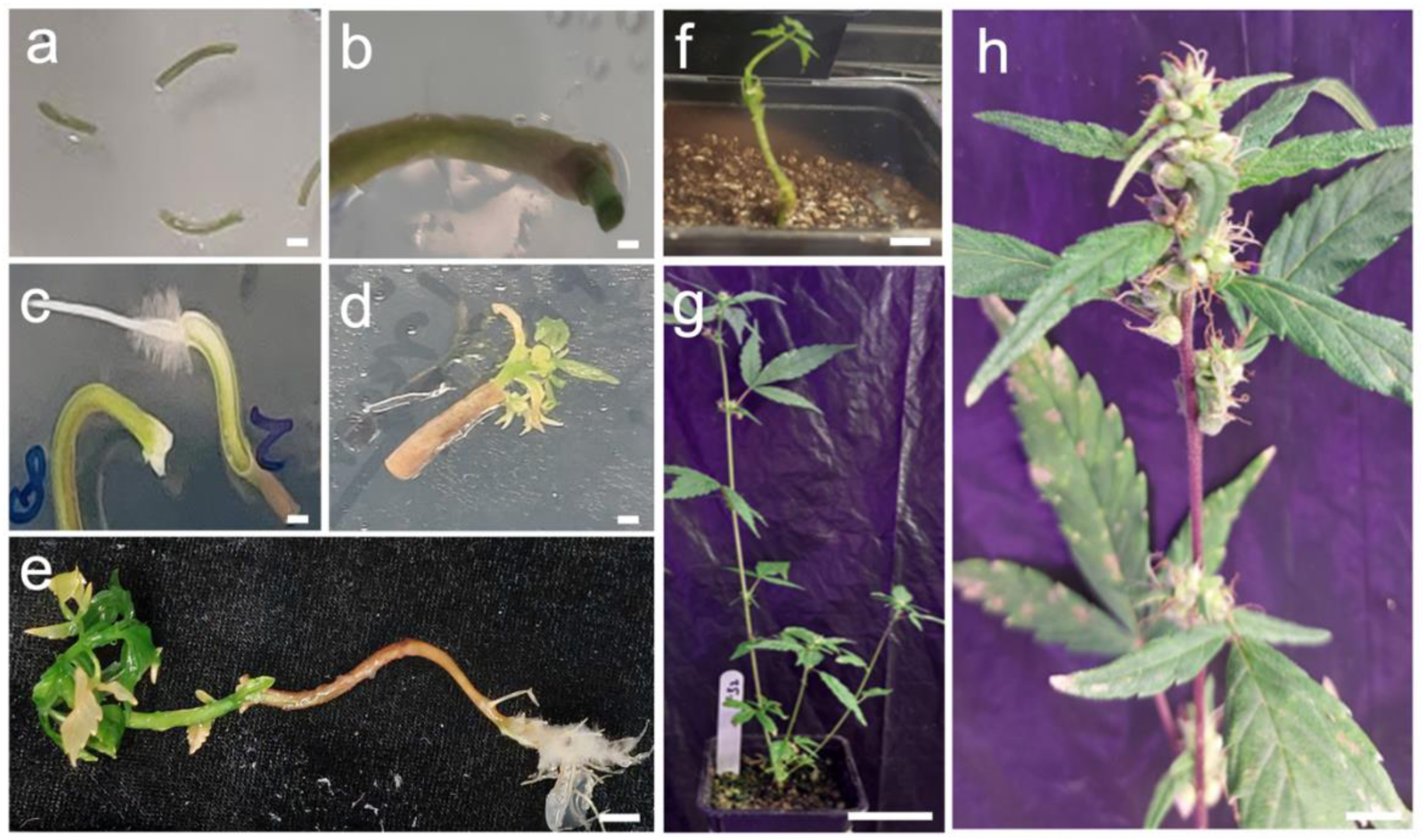
*In vitro* hypocotyl-mediated regeneration and *ex vitro* establishment of hemp. Hypocotyl explants of the hemp cultivar ’Felina 32’ were excised from aseptically germinated seedlings and inoculated on hormone free growth medium (a). After approximately four days growth of ground tissue was observed (b). Adventitious roots emerged after about five days directly from the hypocotyl explant (c), demonstrating direct root regeneration without an intervening callus phase. Multiple shoots with leaves emerged from the regenerating hypocotyl explant after approximately 7-15 days (d). Both shoots and roots were present on a proportion of explants after a minimum of 25 days (e). Rooted plantlets with shoots were transferred to soil and acclimatized in a vermiculite-soil based substrate under controlled humidity and light conditions (f). Hemp plants reached full maturity and reproductive stage after circa 115 days (g) and were able to produce mature seeds after a total of 135 days of cultivation (h, white arrowhead). Scale bars: (a, e) = 10 mm; (b–d) = 5 mm; (f) = 10 mm; (g) = 50 mm; (h) = 20 mm.

## Discussion

Establishing axenic seedling cultures from surface-sterilised hemp seeds is a prerequisite for hypocotyl-based regeneration and any downstream genetic transformation pipeline, yet this upstream step has received surprisingly little systematic attention in the hemp tissue culture literature, where usually standard sterilisation methods are used (Huang et al., 2011; Yildiz and Er, 2002). Here we present a comparative evaluation of sterilisation strategies for hemp, and examine how treatment chemistry, seed provenance and cultivar influence the subsequent morphogenic competence of hypocotyl explants.

Water, salt and vinegar controls produced near-universal contamination or germination failure across all accessions, confirming that more stringent treatment is a prerequisite for axenic hemp seed culture. Among the treatments evaluated, a 75% ethanol baseline combined with sequential 1% H₂O₂ incubation provided acceptable sterile seedling yields for in-house seed material and performed broadly comparably to NaOCl-based protocols supplemented with Octave or PPM^TM^. The absence of statistically significant differences among the NaOCl-based treatments for in-house seeds suggests that, when contamination load is low, the choice of supplementary biocidal agent is less critical. Here, microbial load rather than sterilant stringency appears to be the primary limiting factor for protocol success.

A consistent finding across the entire dataset was the effect of seed provenance. In-house glasshouse-produced seeds achieved sterile seedling rates approximately four times higher than commercially sourced material, and this difference was statistically significant and robust across accessions and treatment conditions. Commercially sourced seeds showed an average of 10% sterile seedling yield, but supplementation with Octave or PPM^TM^ did lower contamination in some cases. While PPM is an established sterilisation agent in plant cell culture (George and Tripepi, 2001), Octave is mostly used in industrial contexts as a fungizide (Denner et al., 1997; Henricot and Wedgwood, 2013) and is not typically used in plant cell culture. Here we demonstrate that it can be used for surface seed sterilisation although further research regarding its effectiveness in hemp and possible applications for other species is needed. The provenance-driven difference most plausibly reflects a higher and more variable microbial burden in field-produced, commercially or long-term stored hemp seeds relative to fresh seed material produced under controlled glasshouse conditions. Storage duration and conditions likely contribute independently, and batch-to-batch variation within seed sources was considerable throughout our experiments. The practical implication is direct: fresh, in-house produced seeds are strongly preferable for tissue culture work, and protocols benchmarked on laboratory-produced material should not be assumed to transfer to commercially sourced batches without re-validation.

With the developed protocol, we consistently were able to regenerate fertile hemp plants from hemp hypocotyl explants without hormone supplementation, confirming previously published observations (Long et al., 2022). Hemp is unusual in this hormone-free regeneration as approaches in other plant species usually require one or multiple plant growth regulators such as cytokinins or auxins for regeneration (Long et al., 2022). Notably, we observed bipolar regeneration from hypocotyl explants, with shoots and roots developing from opposite ends of the same hypocotyl, which was previously reported as a side effect of hormone-free hypocotyl regeneration (Galán-Ávila et al., 2020). This observation suggests that a polarity remains within the hypocotyl and that cells within the hypocotyl retain totipotency, however, further research detailing the concrete mechanisms and possible specifics in hemp are required to complete our understanding of this process. The consistently high proportion of root-only regenerating explants across conditions suggests that supplementation with cytokinin warrants evaluation as a means of promoting hemp shoot organogenesis. The complete developmental cycle from sterilized hypocotyl explant through to fertile adult plant and viable seed was achieved, confirming that the sterilisation conditions described here do not impair subsequent development or reproductive capacity. Further, we demonstrate that explantation and rooting of the newly generated shoot is not a required step in regeneration of hemp, which potentially lowers handling time and associated contamination risk. We also did not observe callus formation from hypocotyl explants under the described conditions, which is usually achieved by adding plant growth regulators (Ahsan et al., 2024; Hesami and Jones, 2021). For many applications, callus formation is not desirable and plant regeneration from hemp calli can be challenging (Ingvardsen and Brinch-Pedersen, 2023; Movahedi et al., 2016), hence hormone-free regeneration might be a viable route in these cases.

Beyond their effect on contamination, sterilisation treatments appeared to have a marked and independent influence on the morphogenic fate of hypocotyl explants. For in-house generated seeds of the hemp cultivar ’Felina 32’, explants from seedlings previously treated with NaOCl combined with either Octave, or both Octave and PPM^TM^, showed significantly higher rates of shoot and root co-regeneration on hormone-free medium compared to H₂O₂ or NaOCl alone. PPM^TM^ supplementation also significantly reduced explant mortality relative to NaOCl alone. These effects on the morphogenic response are unlikely to be explained solely by differences in explant viability following sterilisation, and the question if there is a mechanistic basis for the apparent pro-regenerative effect of Octave and PPM^TM^ warrants further investigation. Notably, high concentrations of PPM^TM^ have been reported as being phytotoxic and reducing regeneration rates (George and Tripepi, 2001), therefore optimising for the lowest required concentration might warrant further studies. For commercially sourced seeds of the cultivar ’Ferimon’, the pattern was reversed, with H₂O₂-treated explants showing the best regeneration outcomes. Hence, the relationship between sterilisation treatment and morphogenic response could be source or even cultivar-dependent and should not be generalised from a single accession. Therefore, we recommend testing at least two different sterilisation protocols when establishing the method for the first time in any given laboratory environment.

Plant regeneration from hypocotyl explants was achievable across all four accessions examined, demonstrating that this developmental capacity is broadly distributed within industrial hemp. Regeneration frequency and consistency varied substantially among accessions. ’Fedora 17’ and ’USO31’ showed the most favourable regeneration profiles, confirming previous findings (Galán-Ávila et al., 2020), with approximately one third of ’Fedora 17’ explants achieving shoot and root co-regeneration, while ’Kompolti’ and ’Felina 32’ showed significantly lower rates of full regeneration under the same conditions. All accessions retained a high proportion of non-regenerating explants, and inter-accession variation in morphogenic trajectory is expected given the high heterozygosity and phenotypic diversity of industrial hemp (Kavanagh et al., 2025; Trubanová et al., 2026; Vergara et al., 2021, 2017). Protocol optimisation will therefore need to be at minimum accession-specific, and possibly batch-specific.

Taken together, these results support the following practical recommendations. Fresh, in-house produced seeds should be used wherever possible. Where commercially sourced material must be used, sterilisation treatment with NaOCl combined with PPM^TM^ and Octave should be considered. The protocol described here (**Figure 5**) provides a practical and documented starting framework that should be adapted based on seed source, batch history and intended downstream application.

**Figure 5.**
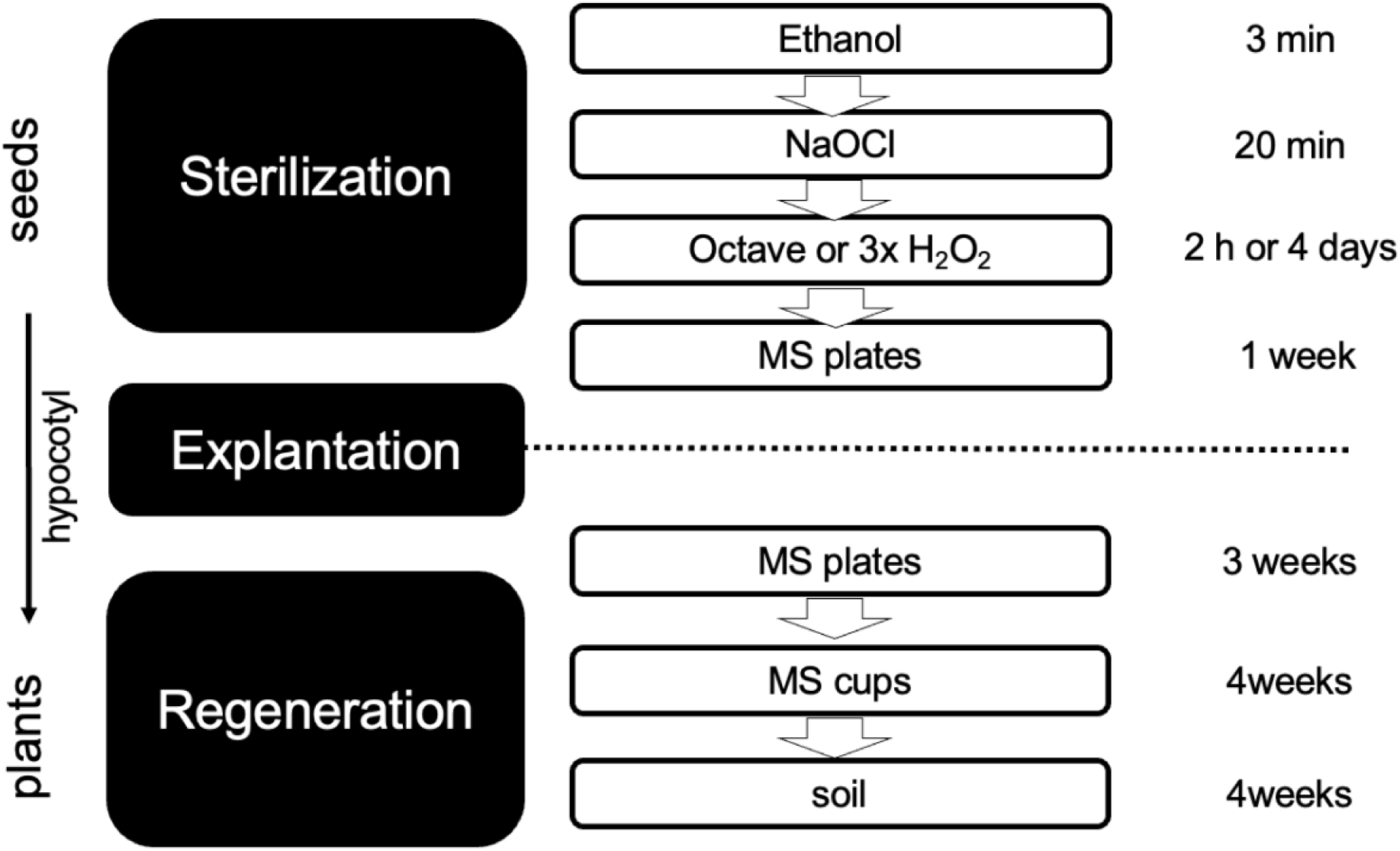
Protocol for sterilisation, explantation and regeneration of hemp (*C. sativa*). The workflow describes sterilisation, explantation and regeneration from seed to seed via hypocotyl explants and regeneration.

## Author Contributions

R.G., S.S., R.M., and A.N. contributed to funding acquisition. S.S., R.M., and J.K. contributed to conceptualization and writing (review and editing). S.S. also wrote the original draft. S.S., R.M., A.D., and A.N. contributed to supervision. R.M. was additionally responsible for project administration. S.S. contributed to data curation. J.K. and G.P. contributed to the methodology. R.G. contributed to visualization and analysis. All authors read and approved the submitted version.

## Acknowledgements

This publication has emanated from research conducted with the financial support of Research Ireland under the Government of Ireland Postgraduate Program Enterprise Partnership Scheme (Grant No. EPSPG/2023/1652). The authors gratefully acknowledge the support of enterprise partner Plantik Biosciences, France. The authors also wish to thank Frances (School of Biological and Environmental Sciences, University College Dublin, Ireland) and Gordon (UCD Rosemount Environmental Research Station) for their valuable technical assistance.

## Supplementary Material

### Supplementary Table

**Table S1. Sterilisation treatment outcomes for in-house and commercial seeds.**

### Supplementary Figures

**Figure S1. Surface sterilisation efficacy of chemical and baseline treatments on hemp seeds.**

**Figure S2. Effect of NaOCl-based sterilisation treatments on hemp seeds of the cultivar ’Ferimon’.**

**Figure S3. Effect of different sterilisation additives on the regeneration outcomes** and viability of hemp hypocotyl explants of the cultivar ’Ferimon’.

