## supplementary_figures for "The Viability Gambit: An Optimized Sterilisation Protocol for Industrial Hemp (*Cannabis sativa* L.) Seeds": Supplementary_Figures_Sterilization_Manuscript copy.pdf

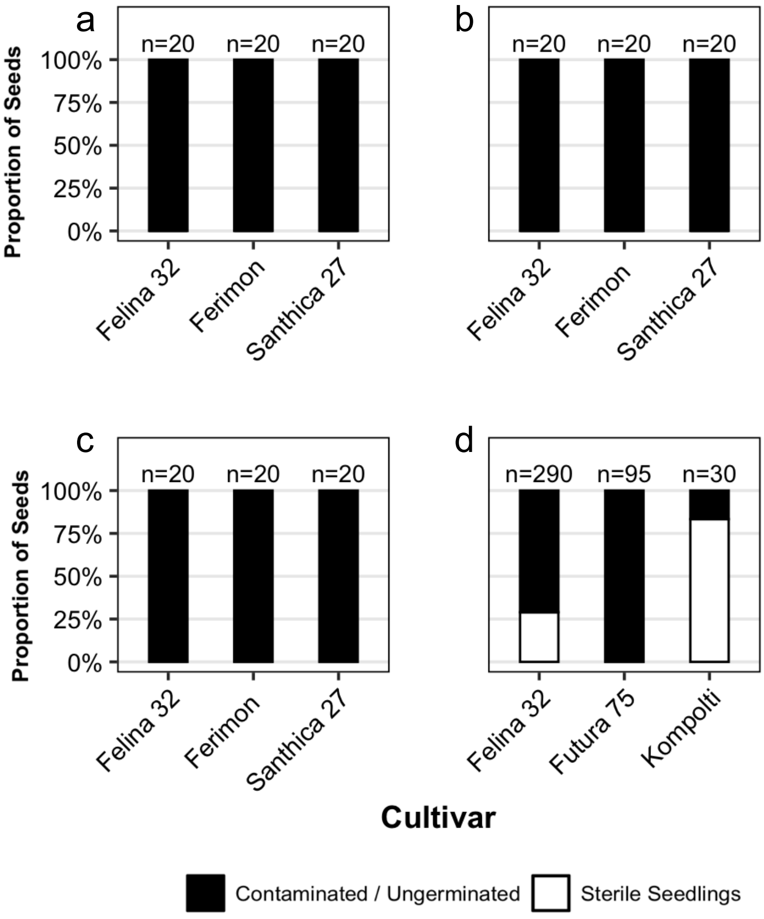


**Figure S1. Surface sterilisation efficacy of chemical and baseline treatments on hemp seeds.** Water control (a) and basic treatments of seeds with salt (b) and vinegar (c) did not yield any sterile seedlings across all evaluated hemp cultivars.Treatment with 1% $H_2O_2$ successfully surface-sterilized seeds (d), though its efficacy was strongly influenced by cultivar (Fisher's exact test, ***, p<0.001). Stacked bar charts display the proportion of seeds yielding sterile seedlings (white bars) or exhibiting contamination/germination failure (black bars). Total sample sizes (n) contamination/germination failure (black bars). Total sample sizes (n) are displayed above each bar.

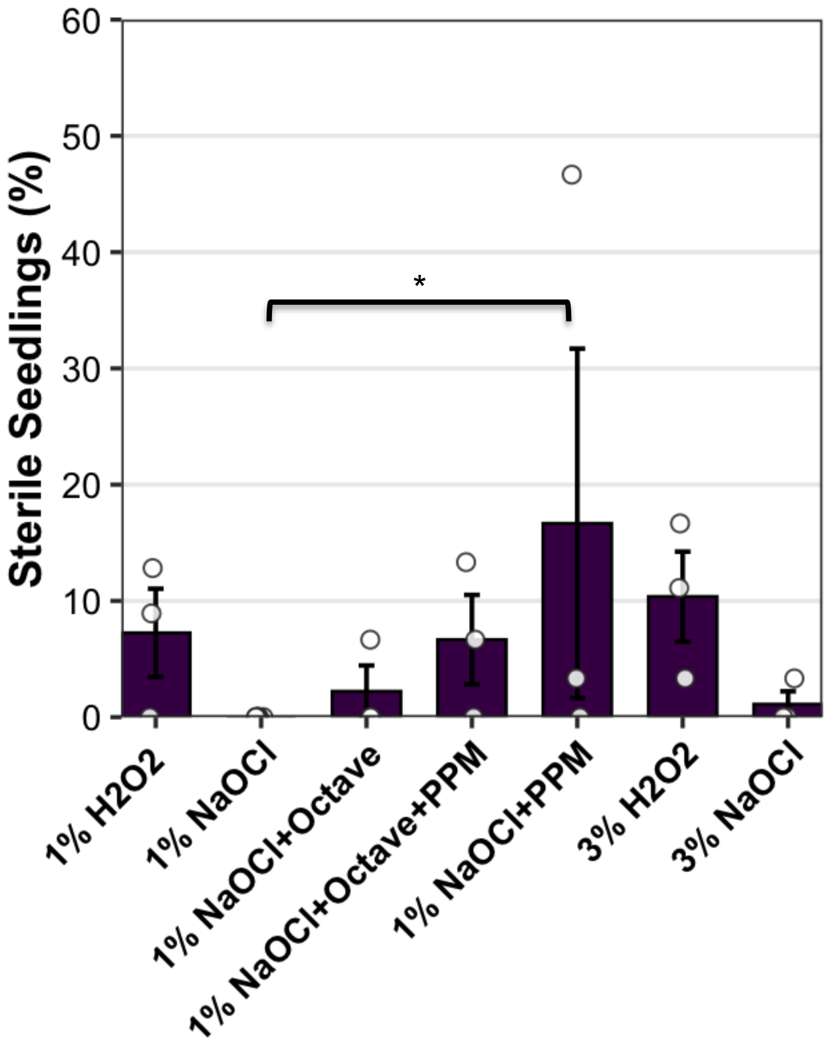


**Figure S2. Effect of NaOCl-based sterilisation treatments on hemp seeds of the cultivar 'Ferimon'.** Commercial hemp seeds of the cultivar 'Ferimon' were treated with different NaOCl sterilisation protocols (1% NaOCl (n=184), 1% NaOCl and Octave (n=89), 1% NaOCl and PPM$^{TM}$ (n=90), 1% NaOCl and Octave and PPM$^{TM}$ (n=90), 1% $H_2O_2$ (n=125), 3% $H_2O_2$ (n=114), and 3% NaOCl (n=96)) and scored for contamination. Significant difference indicated with brackets (Tuckey's Pairwise comparison, *, $p<0.05$). Three independent biological replicates (n=3). Error bars indicate standard error.

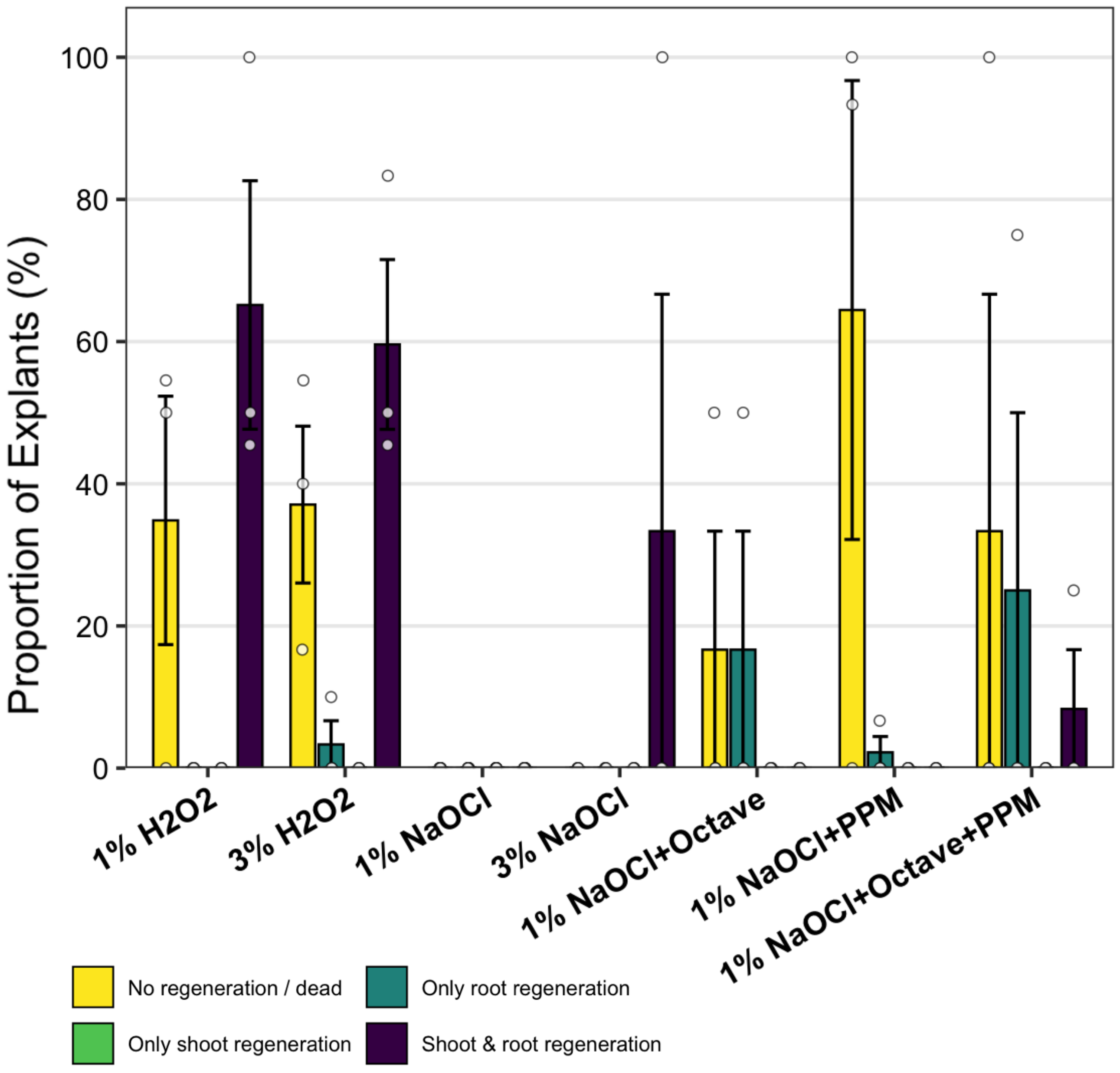


**Figure S3. Effect of different sterilisation additives on the regeneration outcomes and viability of hemp hypocotyl explants of the cultivar 'Ferimon'.** Explants derived from seedlings of the hemp cultivar 'Ferimon' which underwent different sterilisation protocols (1% NaOCl (n=0), 1% NaOCl and Octave (n=2), 1% NaOCl and PPM™ (n=16), 1% NaOCl and Octave and PPM™ (n=5), 1% $H_2O_2$ (n=26), 3% $H_2O_2$ (n=27), and 3% NaOCl (n=7)) were scored for their regeneration and categorized as no regeneration (yellow), only root regeneration (teal), only shoot regeneration (green) and shoot and root regeneration (dark blue).
